# Unifying design principles of endocrine gland mass and its regulatory circuits

**DOI:** 10.1101/2023.07.03.547486

**Authors:** Moriya Raz, Tomer Milo, Yael Korem Kohanim, Omer Karin, Avichai Tendler, Alon Bar, David S. Glass, Avi Mayo, Uri Alon

## Abstract

Hormones are regulatory molecules that impact physiological functions. Much is known about individual hormones, but general rules that connect the regulatory logic of different hormone systems are limited. In this study, we analyzed a range of human hormone systems using a mathematical approach to integrate knowledge on endocrine cells, target tissues and regulation, to uncover unifying principles and regulatory circuits. We find that the number of cells in an endocrine gland is proportional to the number of cells in its target tissues, as one single endocrine cell serves approximately 2000 target cells. We identified five classes of regulatory circuits, each has specific regulatory functions such as homeostasis or allostasis. The most complex class includes an intermediate gland, the pituitary, which can otherwise be considered redundant and exposes to fragilities. We suggest a tradeoff: with the price of fragilities comes advantages -amplification, buffering of hypersecreting tumors, and faster response times. By elucidating these unifying principles and circuits, this study deepens our understanding of the control of endocrine processes and builds the foundation for systems endocrinology.

## Introduction

Hormones are crucial regulatory molecules secreted by endocrine cells into the circulation where they affect target tissues(Melmed *et al*, 2019). Hormones control many physiological functions. Typically, they are produced by dedicated glands whose cells specialize in making the hormone according to specific signals. The hormones flow in the circulation and reach distant tissues, where they are sensed by receptors. Hormones can trigger metabolic, developmental and behavioral effects, and are major factors that affect physiology. Dysregulation of hormones underlies a wide range of pathologies, including diabetes, reproductive disorders and mood disorders(Molina, 2006).

Humans have several specialized endocrine glands, including the hypothalamus and pituitary in the brain, the thyroid and parathyroid at the throat, the adrenal above the kidneys and the ovaries/testes. Hormones are also secreted from the liver, kidney, gut and other organs. Hormones vary in their biochemistry, including peptide, steroid and amino-acid like hormones. The secretory glands vary in size by many orders of magnitude. The physiological effects of the hormones are diverse, ranging from highly specific targets such as thyroid-stimulating hormone (TSH) that has a direct effect mainly on the thyroid gland, to hormones that directly affect virtually the entire body such as cortisol. Some hormones provide homeostasis to key metabolites, whereas others control acute responses to stimuli. Some hormones affect behavior, some affect the immune system and others have systems-level metabolic effects. The endocrine systems also vary in their regulatory logic -some hormones are secreted by neurons, others are secreted under the control of other hormones, and yet others are secreted in response to metabolic signals.

In order to make sense of such diverse aspects of biology, it is useful to define recurring patterns and concepts(Milo *et al*, 2002). These patterns can help guide understanding, and are also crucial to form research hypotheses that take concepts from known systems and translate them into new experiments on lesser explored ones. For example, feedback regulation is a hallmark of many homeostatic systems such as insulin control of glucose. The discovery of such feedback loops led to mathematical models that resulted in formulas that are useful for clinical research -such as estimates for insulin resistance known as the HOMA-IR formula(Duncan *et al*, 1995; Matthews *et al*, 1985). Analogy between systems allows researchers to carry over ideas from one system to another, such as development of HOMA-like formula for the thyroid(Dietrich *et al*, 2018; Chatzitomaris *et al*, 2017), and the discovery of long feedback loops in the hypothalamic-pituitary hormone axes.

More recent work in our group has found principles related to changes in endocrine gland mass -such as the ability of gland mass to grow or shrink in order to compensate for physiological changes such as insulin resistance or chronic stress (Karin *et al*, 2020, 2016), to generate hormone seasonality (Tendler *et al*, 2021) and to explain the transition between subclinical and clinical autoimmune disease in which upstream glands can partially compensate for destruction of tissue (Korem Kohanim *et al*, 2022).

There remain many open questions. For example: what determines the size of different glands? and what determines the regulatory logic of their circuits? It would be important to discover additional unifying principles to deepen our understanding of endocrine systems.

Here, in a search for unifying principles, we study a wide range of human hormone systems using a mathematical approach to integrate knowledge on endocrine cells and their targets and regulation. We find that gland mass is proportional to the mass of the total target tissues, indicating that a single endocrine cell serves about 2000 target cells. We find that diverse systems can be organized into 5 classes of regulatory circuits, each with specific regulatory functions. We show how the pituitary gland, a key element of several endocrine circuits, can offer functional advantages over alternative designs without such an intermediate gland. This includes buffering against hormone-secreting tumors and providing speedup to response to chronic stress. These principles can add to the conceptual basis of systems endocrinology.

## Results

### Endocrine gland size is proportional to the size of its target tissue

Endocrine glands vary widely in size. The glands also differ in the number of cells they serve, in the sense of the number of cells that respond to the hormone. The adrenal and thyroid produce hormones that are sensed by virtually all cells in the body. Other glands are much more specialized.

We asked whether there is a relationship between the number of secretory cells in an endocrine gland, which we call gland size, and the total number of cells in its target tissues. For this purpose we used a literature search for each of 24 hormones (references in SI table 1) to determine the expression of receptors. We included target tissues that express high levels of the receptor and where the hormone has a documented physiological function. In cases where there were multiple estimates of organ mass we use the geometric average (Biology by the numbers, Milo)(Philips). The detailed calculations for each hormone are detailed in the SI.

**Table 1:**
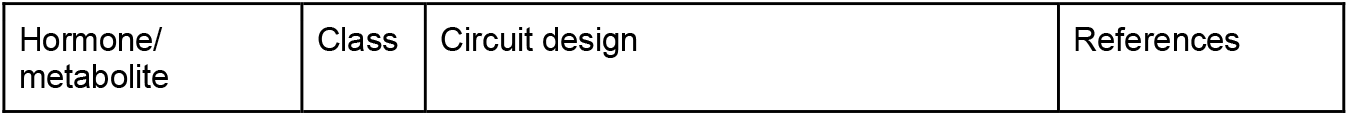
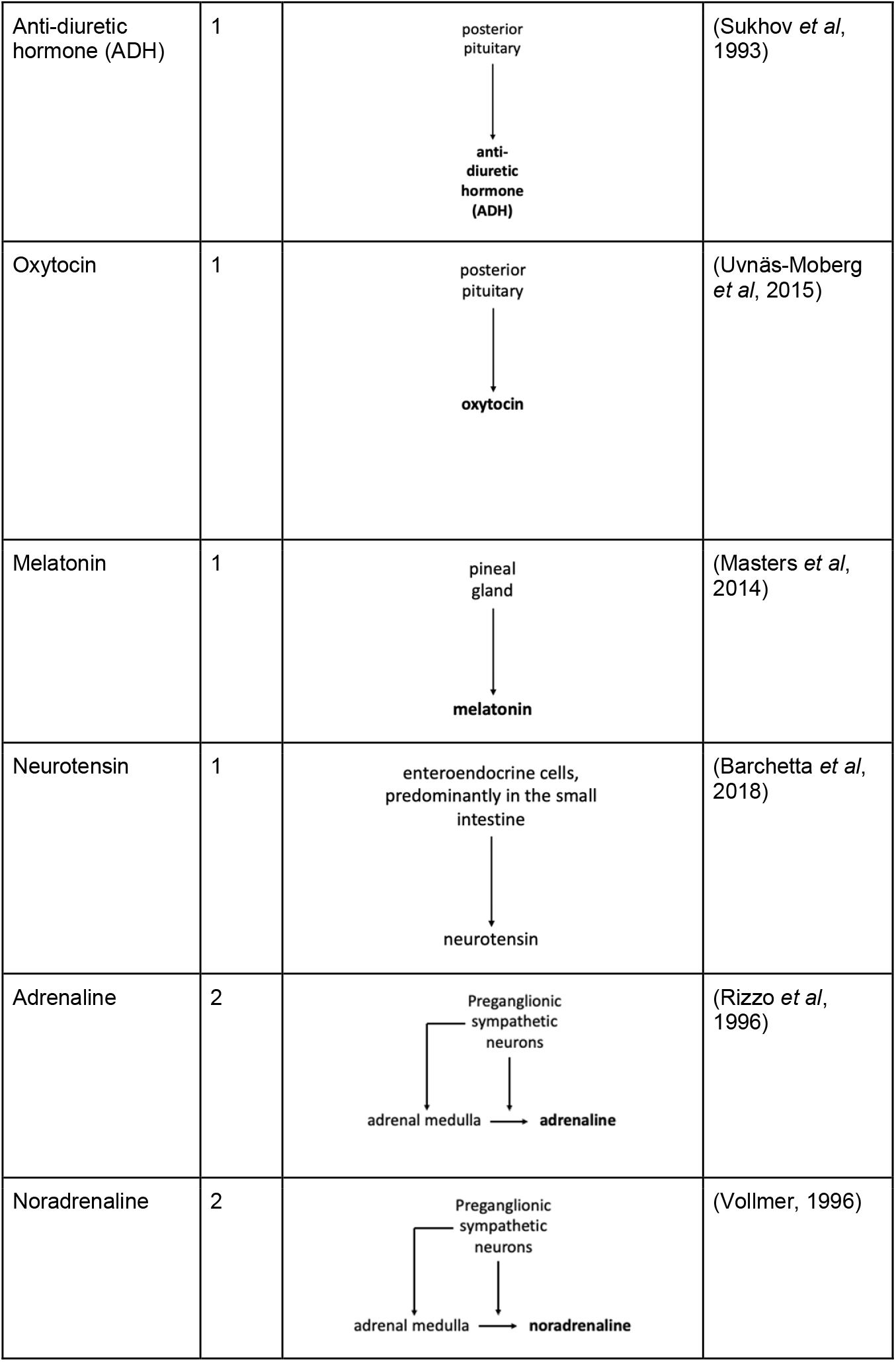
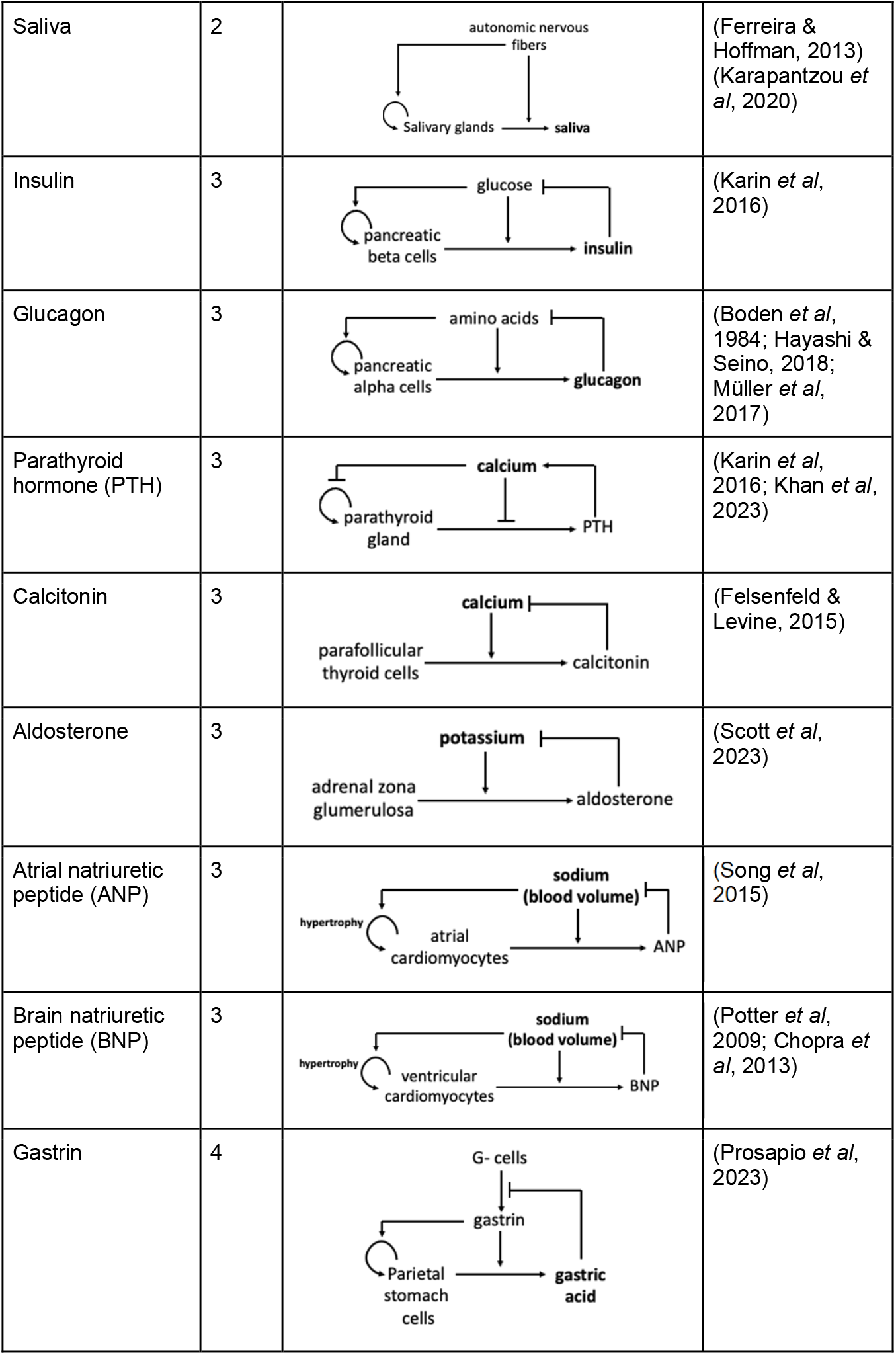
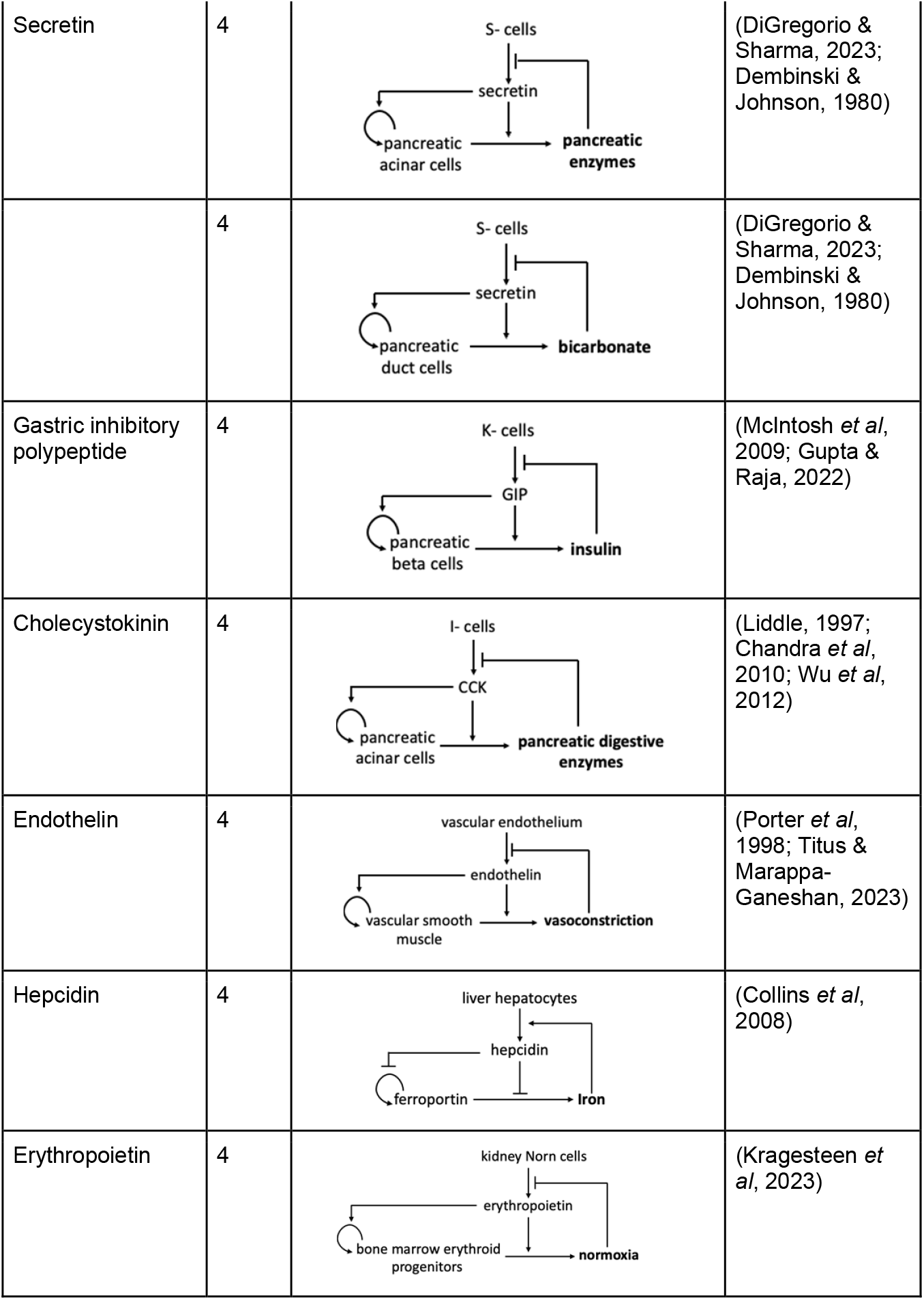
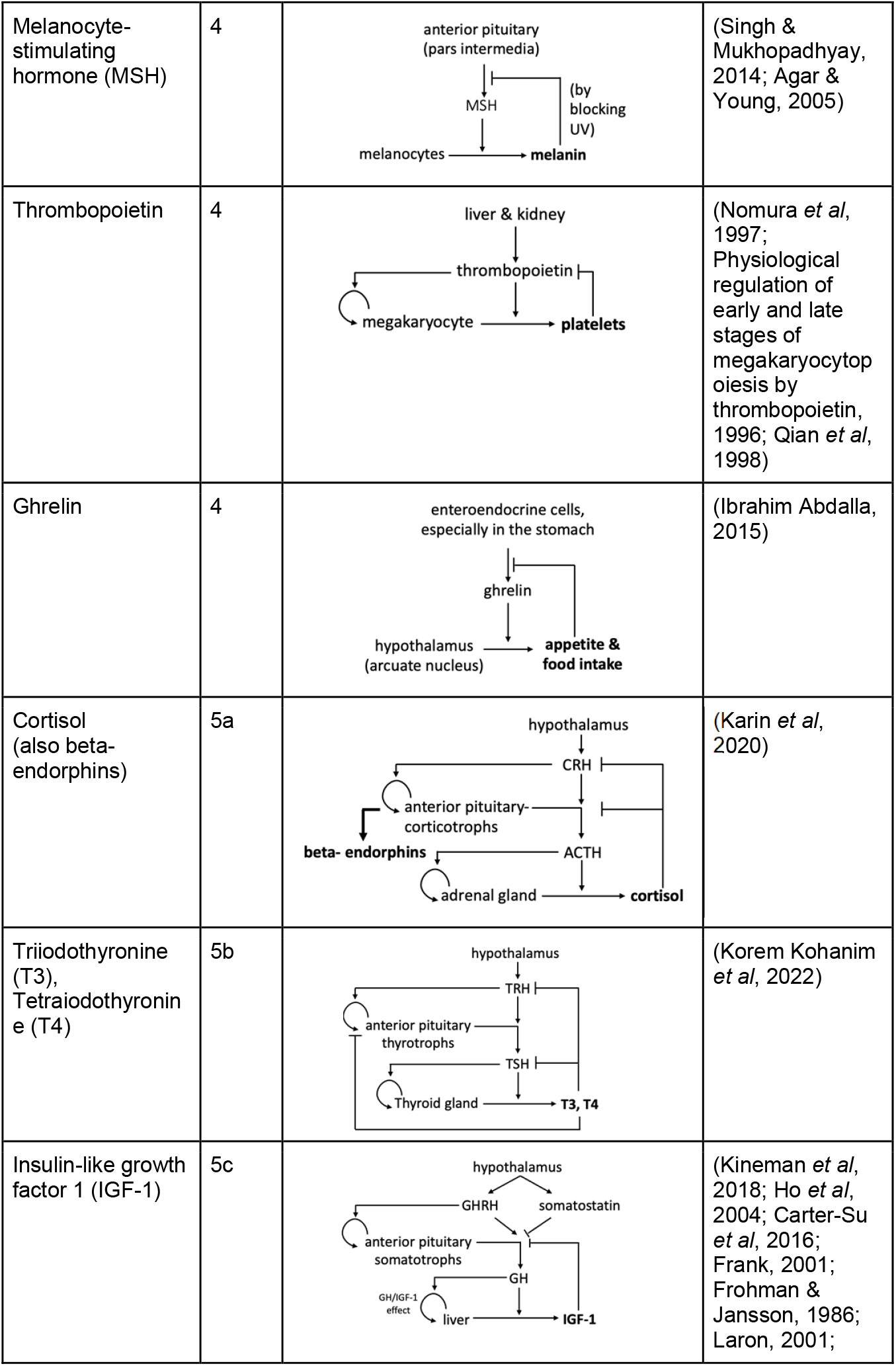
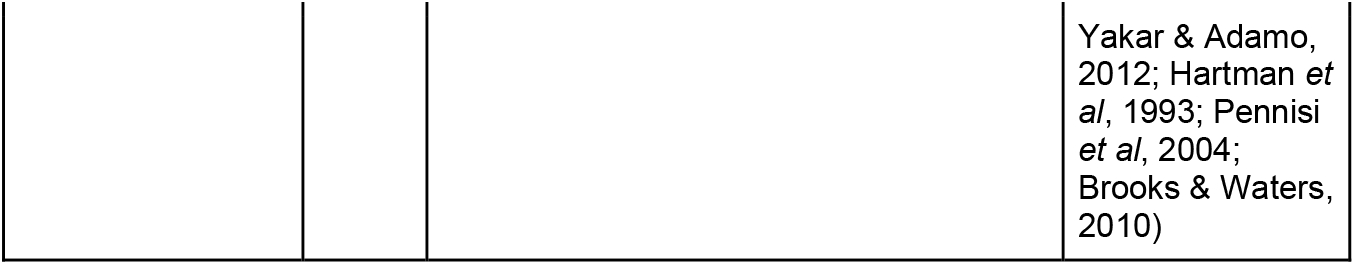
Examples for each endocrine circuit class.

We excluded hormones which are not produced by a dedicated endocrine cell type, but rather are produced by cells that have other major functions. For example, leptin and adiponectin are made by fat cells but the production of these hormones is not the main task of these cells which is storage and metabolism of fat. For the same reason we excluded liver-made hormones including IGF1, hepcidin and thrombopoietin.

We find that target size and gland size are linearly proportional to each other (*R*^2^ = 0.89).The slope of the regression line is close to one (0.95). The ratio of target to secreting cell numbers averages 10^3.3±0.56^. This roughly corresponds to two thousand target cells served by each dedicated endocrine cell.

This proportionality may stem from a maximal hormone production rate per unit biomass. This suggests a principle in which a gland size evolved to serve the size of its target tissues. According to this principle, hormones secreted as a secondary function of a tissue (which has a non-endocrine major function) should lie below the line-the mass of the gland should be much larger than its predicted target size. This is the case for hormones secreted by fat (e.g., leptin which targets the brain and immune system) and liver (e.g., thrombopoietin whose target is platelets and megakaryocytes) which have gland sizes of 10^10^− 10^11^ cells; their targets are much smaller than the 10^13^− 10^14^ cells that would be predicted by the proportionality in Fig 1 (the total number of cells in human is about 3 ∗10^13^ (Sender *et al*, 2016).

**Figure 1:**
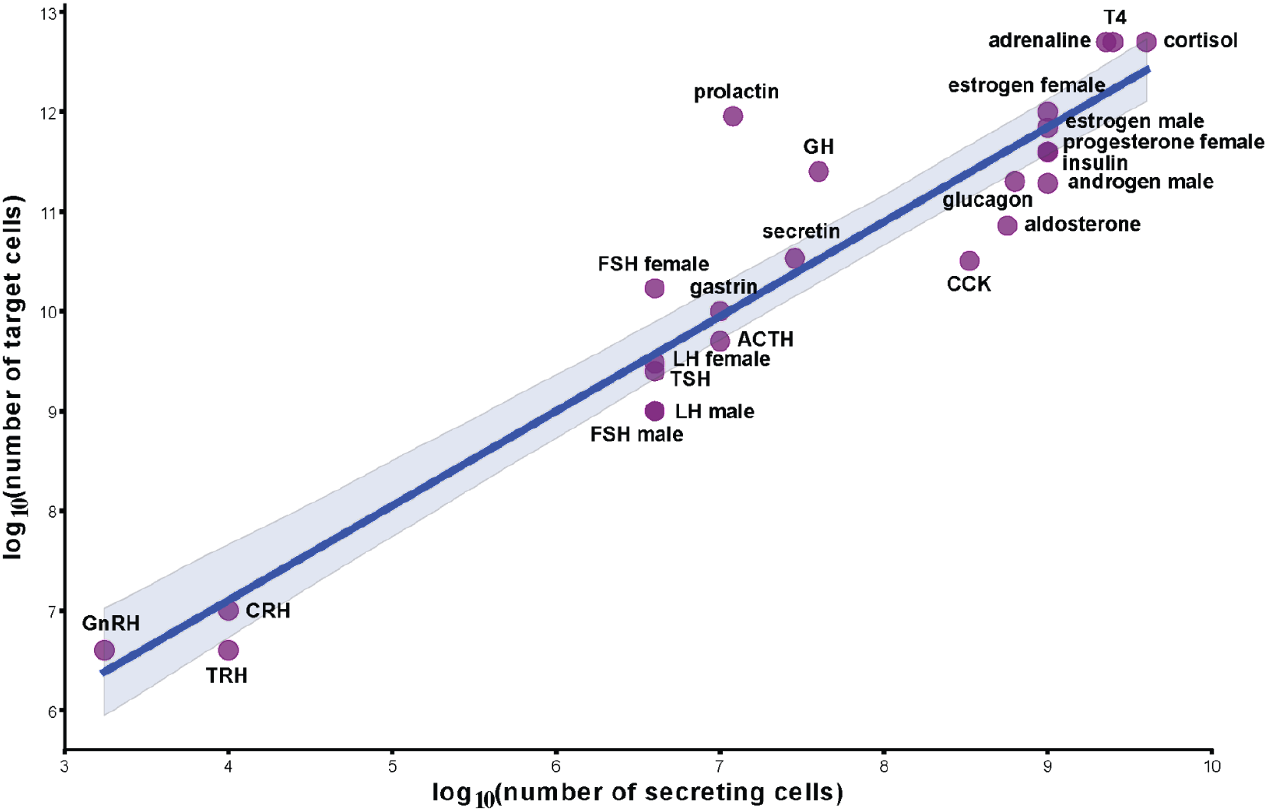
Gland cell number is approximately proportional to total target tissue cell number. Regression line and 95% CI are shown in blue. Error bars on the estimates for the number of cells are about a factor of two, but are not easy to estimate due to sparse literature. (ACTH= Adrenocorticotropic Hormone, CCK= Cholecystokinin, CRH= Corticotropin Releasing Hormone, FSH= Follicle Stimulating Hormone, GH= Growth Hormone, GnRH= Gonadotropin Releasing Hormone, LH= Luteinizing Hormone, TRH= Thyrotropin Releasing Hormone, TSH= Thyroid Stimulating Hormone, T4= Tetraiodothyronine).

### Five classes of hormone circuit motifs serve specific dynamical functions

We next ask about the control of gland sizes in endocrine systems. The data in the previous section concerns the mean gland size, but gland size can often change over time according to physiological conditions.

Control of gland size is interlocked with the feedback control of the hormone levels(Karin *et al*, 2020; Korem Kohanim *et al*, 2022; Karin *et al*, 2016). We thus consider two levels of regulation. The first is regulatory signals that induce the cells of the gland to secrete the hormone. These are neural, metabolic and endocrine signals. This level of regulation works on the timescale of the hormone half-life which ranges from minutes to hours for most hormones (thyroxine T4 is an exception with a half-life of 7 days) (Melmed *et al*, 2019). The second level of regulation is less studied. This is regulation on the functional mass of the gland. Many endocrine glands have growth factors, and in many cases the major growth factor is also the regulatory signal that also instructs the cells to produce and secrete the hormone (Karin *et al*, 2016, 2020; Korem Kohanim *et al*, 2022). The timescale of this level is determined by the turnover time of the endocrine cells, and is on the order of weeks to months.

This principle was recently used to study the dynamics on the scale of weeks of the Hypothalamic-Pituitary-Thyroid (HPT) (Korem Kohanim *et al*, 2022) and Hypothalamic Pituitary Adrenal (HPA) (Karin *et al*, 2020) axes, as well as the pancreatic beta cell circuit (Topp *et al*, 2000; Karin *et al*, 2016; Ha *et al*, 2016).

We performed a literature search to systematically categorize the regulation of 27 hormones at these two levels. When considering these two levels of regulation, we found that all the endocrine systems we studied can be organized into five classes of circuits, which we term Classes 1-5. In each class, the same circuit logic appears in different hormone systems. Each of the five circuit classes can thus be described as a circuit motif. Note that these circuits differ from classical gene-regulatory motifs(Alon, 2007) in including both signaling and cellular growth. Class 1 circuits are the simplest and class 5 are the most complex.

We note that the number of possible regulatory circuits is much larger than 5. For example, if one enumerates all possible connected circuits of three glands one obtains 512 possible circuits (6 interactions between glands and 3 possible autocrine arrows). If one includes all combinations of possible regulatory signs on the arrows (each either negative or positive) this number grows to 3^9^ = 19683 possibilities. Thus it appears that physiology utilizes only a tiny fraction of the possibilities, suggesting meaningful function for the 5 classes.

For each class of circuits we also developed a minimal mathematical model (Box 1). These models describe the dynamics of the hormone concentration, metabolite concentrations and the gland functional mass. We used as the basis the classical Topp model for the insulin system (Topp, 2000) and a mathematical model for the HPA axis by Karin et al (Karin *et al*, 2020). These models include the minimal number of parameters (production rates, removal rates, growth rates) needed to describe the essential mechanism. With this approach we constructed a model for each of the classes, and used it to study its dynamics function.

**Box 1.**
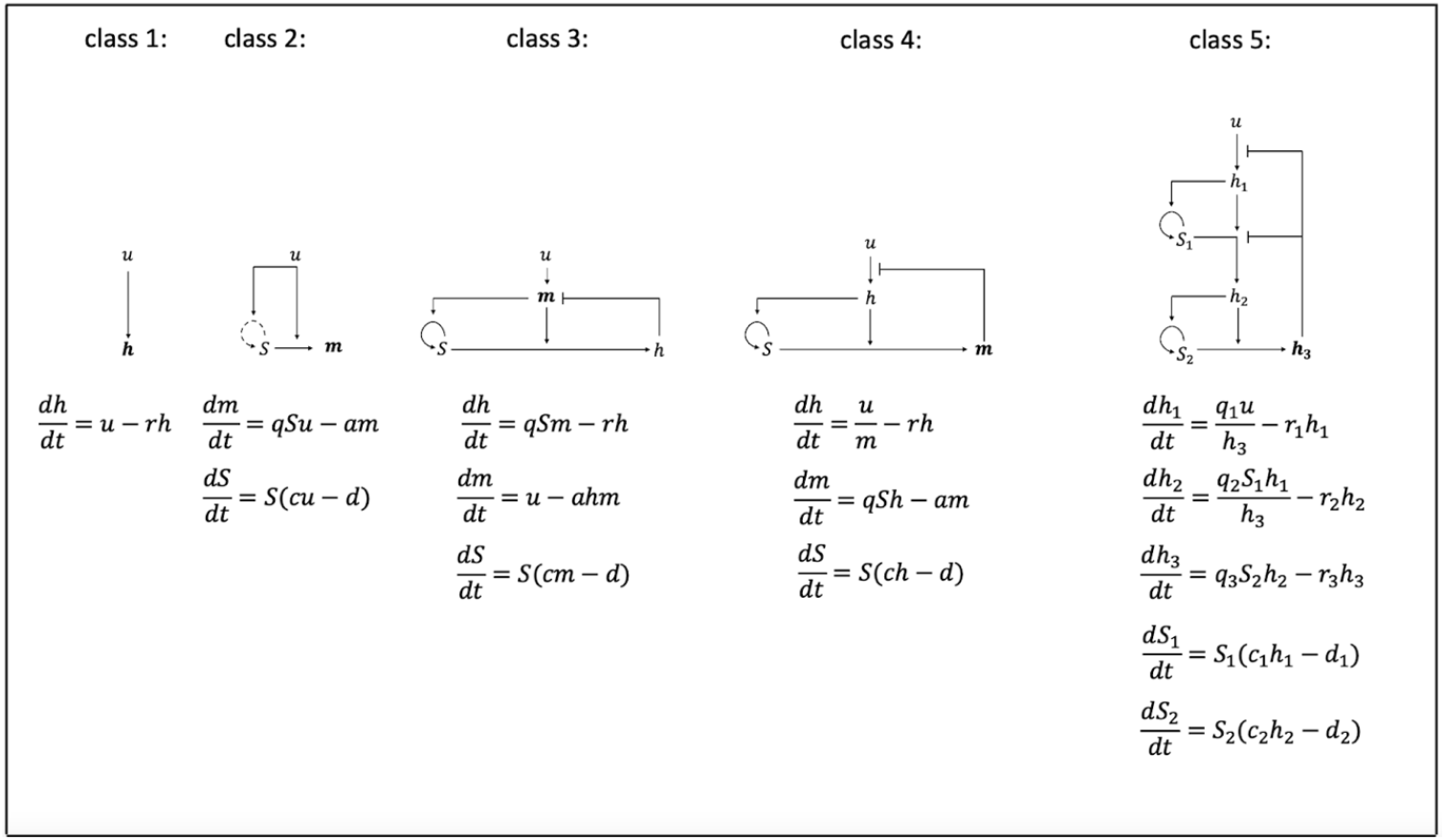
Simplified equations for the five classes of hormone circuits. h= hormone concentration, m=metabolite concentration, S=endocrine gland mass, u=input signal, r=hormone removal rate, q=hormone/metabolite production rate, a=metabolite removal rate, c=cell proliferation/growth rate, d=cell death or removal rate.

Class 1 (Figure 2A) circuits are simply neurons that secrete a hormone. An example is the hypothalamic neurons that secrete Antidiuretic hormone (ADH) and oxytocin. Class 2 (Figure 2B) circuits have a neuronal input to an endocrine or secretory cell, with or without instruction for cell hypertrophy or hyperplasia. Examples include sympathetic control of adrenal medulla cells that secrete adrenaline and neuronal control of salivary glands. Both class 1 and class 2 circuits are input-output devices that convert a neuronal input into a secretion rate of a hormone or metabolite.

**Figure 2:**
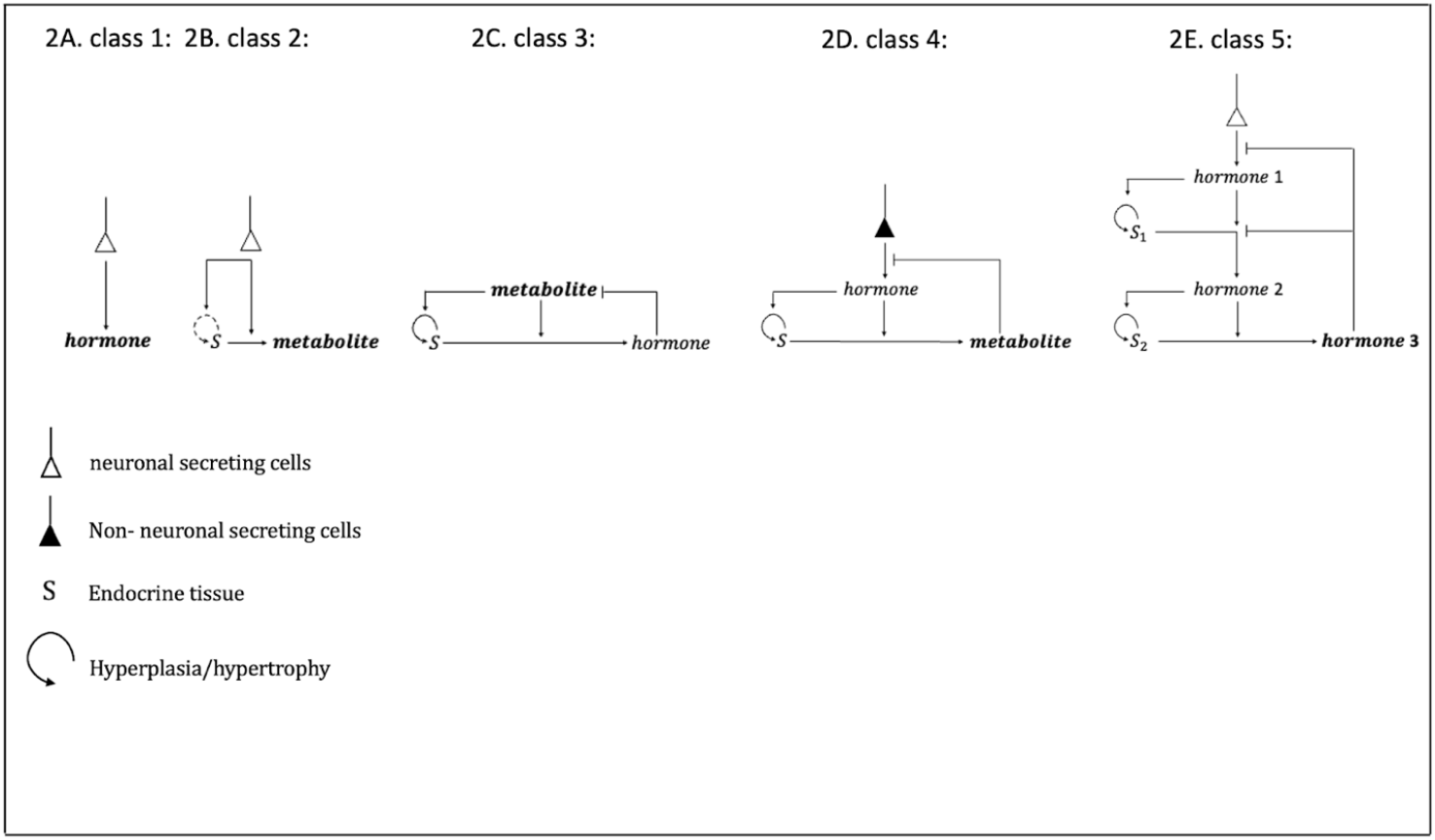
Five classes of hormone circuits and their basic structure.

In class 3 circuits (Figure 2C), endocrine cells secrete a hormone in response to a metabolic signal. The hormone acts to restore the metabolite to a homeostatic level. The metabolite also regulates the endocrine cell growth (hyperplasia or hypertrophy). Examples are beta cells which secrete insulin under control of blood glucose. Glucose is a beta cell growth signal primarily by hypertrophy in humans after childhood (Cerf *et al*, 2012; Jones *et al*, 2010). Another example is the parathyroid chief cells which secrete PTH under control of blood free calcium ions. Free calcium ions also act to regulate parathyroid cell proliferation (Karin *et al*, 2016).

Class 3 circuits, unlike class 1 and 2, can achieve robust homeostasis of their metabolite input signal by means of the following mechanism. The cells respond within minutes to changes in metabolite, such as postprandial insulin secretion. They also respond within weeks by changing their gland functional mass to compensate for physiological changes. As long as the metabolite is away from its set point the cell mass grows or shrinks until the set point is achieved. An example is the hypertrophy of beta cells seen in individuals with insulin resistance. The change in gland mass can compensate precisely for changes in physiological parameters, as long as the gland is not limited by a maximal size (Karin *et al*, 2016).

Class 4 (Figure 2D) circuits describe secretory cells whose input signal is a hormone, rather than a metabolite as in class 3. These cells secrete a hormone or metabolite under control of the input hormone. The input hormone is also a growth factor for the cells. The input hormone is itself secreted by another cell type. For example, stomach parietal cells secrete acid under control of the input hormone gastrin, which itself is secreted by other cells in the digestive tract called G-cells. Gastrin is also a growth factor for the gastric parietal cells.

This circuit can provide allostasis -the output hormone/metabolite have steady state set points that can be tuned to physiological needs. This tuning can be achieved by changing the secretion rate of the input hormone and other parameters. In addition the class 4 circuit locks the input hormone to a homeostatic value on the slow timescale, similar to class 3 circuits.

The difference in function between class 3 and 4 is due to the position of the hormone in the circuit (see also equations in Box 1). The input signal in class 3 is a metabolite, and in class 4 is a hormone. When a signal controls the cell growth rate, it participates in an integral feedback loop that locks the input signal to a fixed point on the scale of weeks. In class 3 circuits, metabolites are thus locked to a constant value, 5mM in the case of glucose. In class 4 circuits, the input hormone is locked but the output, such as stomach acid, depends on the gland mass which can adjust to varying parameters.

Finally, class 5 (Figure 2E) circuits are the most complex. They involve three glands, a top hypothalamic gland and two downstream glands -a pituitary cell type and an effector gland. The pituitary and effector glands can change their functional mass. This circuit resembles two instances of a class 4 circuit placed on top of each other in series. The size of the glands shows a hierarchy where the top neuronal gland is smallest, the middle (pituitary) gland cell type is intermediate and the effector gland is the largest. This is due to the principle of Fig 1 where gland mass is proportional to its target mass. There are several subtypes of class 5 circuits, each with a different pattern of interactions. Class 5 circuits are found in the hypothalamic-pituitary axes. These axes control major functions in vertebrates. The HPA axis controls stress response via the hormone cortisol. The thyroid axis controls metabolic rate via thyroid hormones. Similarly, the sex hormone pathway and growth hormone pathways share a class 5 design.

### The pituitary as an endocrine amplifier

Next, we explore the structure-function relationship of class 5 circuits. One fundamental question is what advantage this design, with a pituitary gland in the middle, might have compared to simpler circuits.

We propose that one function of the pituitary is to act as an amplifier of the hypothalamic hormones.

We saw in Fig 1 above that a single endocrine cell can serve about 2000 target cells on average. In the HP class 5 axes, a tiny brain region, the hypothalamus, secretes hormones. If there were no pituitary, the hypothalamus would need to secrete enough hormones for the effector gland, such as the adrenal or thyroid. These glands in turn serve the entire body and thus have about 10^10^ cells. Without a pituitary, the hypothalamic regions that secrete each hormone would thus need to have a mass of about 10^7^ cells, which is 3 orders of magnitude larger than observed. It may be implausible to host such a large number of cells in the hypothalamus. The pituitary, which lies external to the skull, can more easily host a large number of endocrine cells. It may thus have an amplification role, allowing 10^4^ hypothalamic cells to produce enough hormone for the 10^7^ pituitary cells, which then provides enough hormones for the 10^10^ cell effector gland.

### The pituitary can compensate for toxic adenomas up to a threshold

Beyond this amplification role, we asked whether the pituitary also has dynamical functions. We begin by noting that changes in the pituitary mass can buffer physiological and pathological variations. An example has been described previously in the context of thyroid disease (Korem Kohanim *et al*, 2022).

Here we add to this previous work by studying the ability to compensate for tumors that hypersecrete hormones in the HPA axis. These tumors arise quite frequently. Known as incidentalomas, they are found in up to a few percent of individuals(Jing *et al*, 2022). The tumors usually have no physiological consequence-the hormone levels are normal. When the tumors exceed a threshold size, they dysregulate the hormone levels, causing overt hypercortisolism called Cushing’s disease.

We asked what sets the threshold between subclinical and clinical disease. We also asked whether there are qualitative differences in the dynamics between tumors in the pituitary and tumors in the adrenal that have the same net effect on cortisol.

We thus analytically solved the HPA mathematical model. We begin with an adrenal tumor that secretes cortisol, Fig 3A. We assumed that the tumor secretion rate is not regulated by ACTH, as commonly observed(Sakai *et al*, 1993). We find that, as long as the secretion rate beta is below a critical threshold, the adrenal mass shrinks to precisely compensate for the extra hormone secreted by the tumor (Fig 3A-D). However, at a critical secretion rate β, the native (non-tumorous) adrenal mass shrinks to zero at steady state (a transcritical bifurcation). Thereafter, as the tumor secretion rate grows, cortisol levels exceed normal levels (Fig 3A-D) and clinical symptoms of hypercortisolism occur, called Cushing’s syndrome.

**Figure 3:**
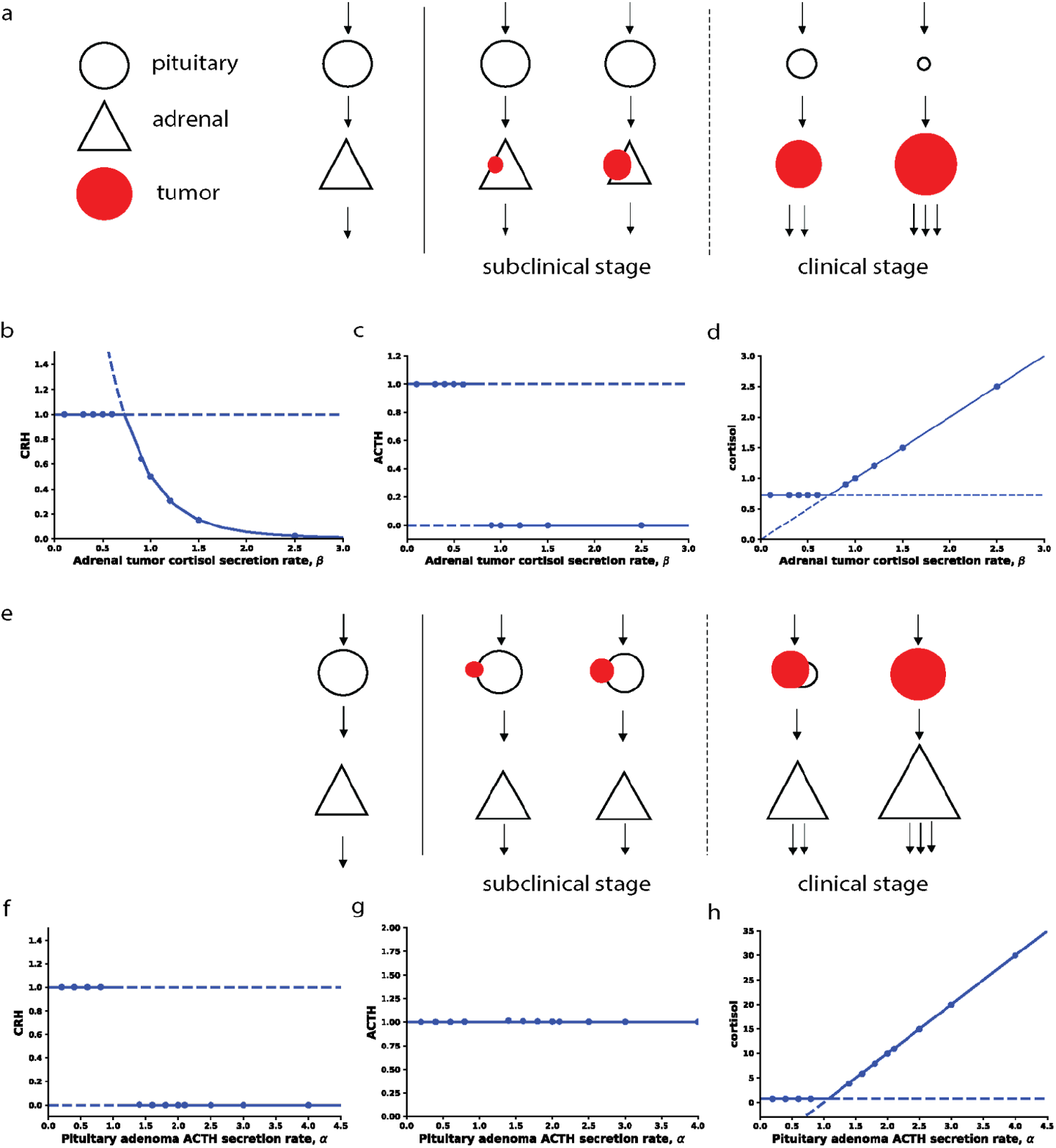
Transition between subclinical and clinical Cushing’s syndrome in the cases of pituitary and adrenal tumors. A) Schematic showing how an adrenal tumor causes the native adrenal mass to shrink. Clinical disease begins when it shrinks to zero. B) CRH C) ACTH and D) cortisol as a function of adrenal tumor cortisol secretion rate beta. E) Schematic showing how a pituitary ACTH-secreting tumor causes the native pituitary corticotroph mass to shrink. Clinical diseases begin when it shrinks to zero. F) CRH G) ACTH and H) cortisol as a function of pituitary tumor ACTH secretion rate alpha.

We compared this adrenal tumor to a different form of Cushing’s pathology in which a tumor in the pituitary secretes ACTH at rate □. We find that for a range of low secretion rates the native (non-tumor) pituitary corticotroph mass shrinks to compensate and maintain a normal level of ACTH and cortisol (Fig3E-H). However, at a critical tumor secretion rate □, the non-tumor pituitary corticotroph functional mass shrinks to zero at steady state. Thereafter, higher tumor secretion causes higher than normal levels of cortisol, resulting in Cushing’s disease (Fig 3E-H). The pituitary case is about 10 times more common than the adrenal case.

In both cases, the system undergoes a transition which, in the language of dynamical systems, is a transcritical bifurcation(Strogatz, 2019). Beyond the transition, the compensating gland functional mass drops to zero.

We conclude that changes in gland mass can compensate for toxic adenomas until they reach a critical mass.

### The pituitary can speed responses on the scale of weeks compared to simpler circuits

We also asked whether the pituitary can provide dynamical benefits to class 5 circuits as compared to simpler circuits. For this purpose we studied the HPA model with prolonged stress inputs that rise and remain high for months and then fall, to explore the onset and recovery from such prolonged stress.

We compared the natural class 5 circuit with two hypothetical simpler designs for cortisol control. One has a pituitary but the pituitary does not change mass-it has a fixed mass. The second is a class 4 circuit without a pituitary. Here the adrenal is directly activated by a hormone from the hypothalamus. To allow a ‘mathematically controlled comparison’(Savageau, 1976; Alon, 2019; Adler *et al*, 2017) we set all hormone half-lives and production rates to be equal between the circuits.

In the natural class 5 circuit, the onset of stress causes a rapid response on the scale of hours, and then a slower increase on the scale of weeks as gland masses change(Karin *et al*, 2020). Similarly, at the end of the stress input pulse, there is a rapid reduction on the scale of hours, followed by a slower adjustment due to gland mass changes on the scale of weeks.

The alternative class 5 circuit with constant-mass pituitary has a slower rise time, defined as the time to first reach 90% of the hormone steady state value (Fig 4). The alternative class 4 circuit has the slowest rise time. The same is found upon recovery from the stress pulse.

**Figure 4:**
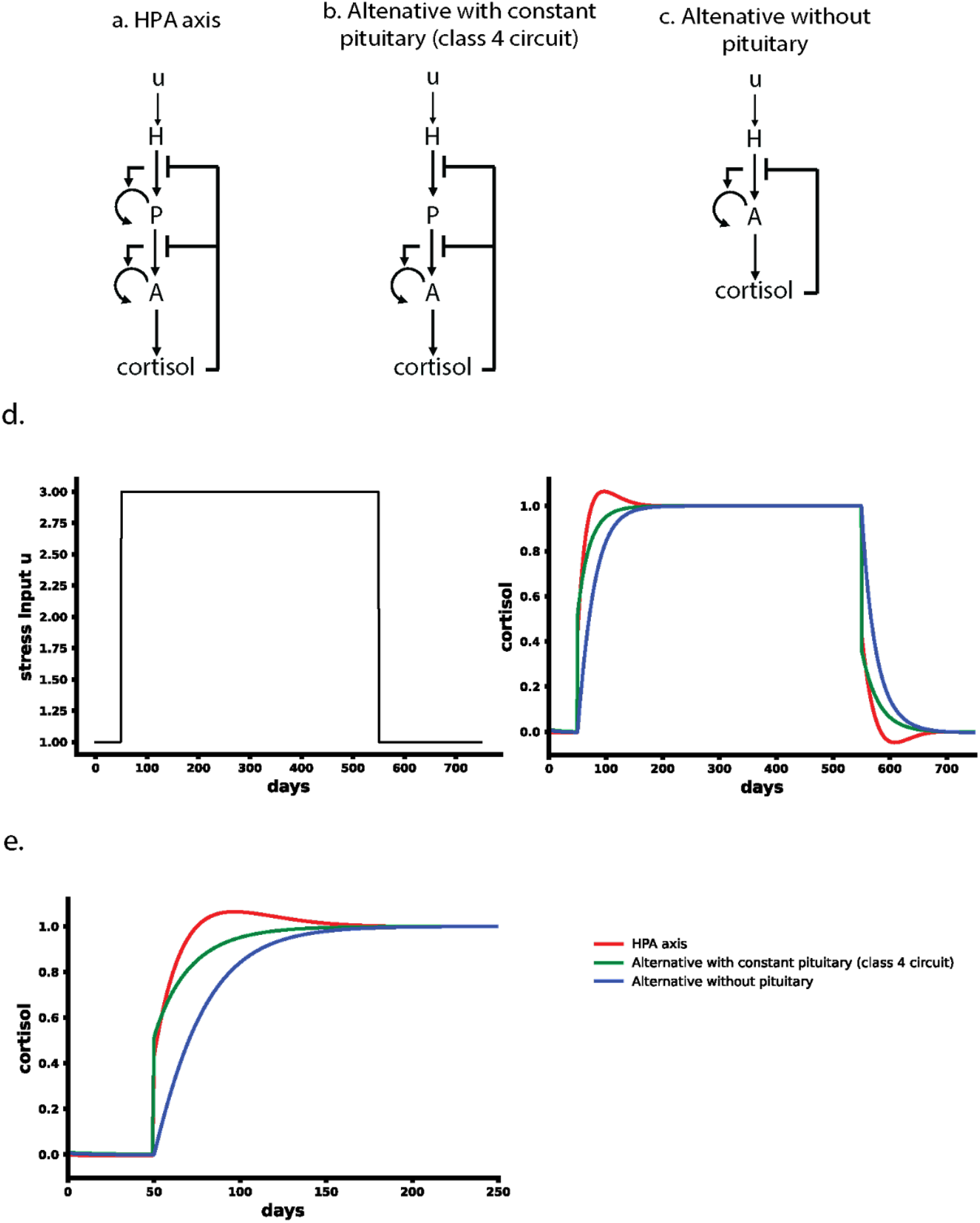
Comparison of alternative HPA designs indicates that the natural design overshoots and has the fastest rise time. A) natural class 5 design of the HPA axis in which the pituitary functional mass is regulated, B) alternative design with a pituitary that has a constant functional mass C) alternative design without a pituitary no middle gland). D) Dynamics of cortisol in response to a prolonged stress input which rises to 3 times the normal input and drops back 500 days later. E) Zoom in on the rise phase shows that the class 5 circuit reaches 90% of the response fastest.

The class 5 circuit also shows an overshoot due to the adjustment of pituitary mass absent from the other two simpler circuits.

We conclude that the pituitary with changing mass can provide a speedup on the scale of weeks when stress conditions change.

## Discussion

We present several design principles for hormone circuits. The mass of each endocrine gland is approximately proportional to the mass of its target tissues. Thus, across different hormone systems, each endocrine cell serves about 2000 target cells. We further find that endocrine systems can be organized into 5 classes of recurring regulatory circuit motifs. In these circuits, gland functional mass can adjust on the timescale of weeks in order to compensate for physiological and pathological changes. Focusing on the HPA axis, we ask about the role of the pituitary gland. This gland might be considered superfluous. We show that it has several important functions. It serves as an endocrine amplifier to supply enough hormone for the large effector gland. We demonstrate how the pituitary gland can compensate for hormone-secreting tumors in the adrenal axis, and how this compensation breaks down at a critical tumor secretion rate to explain the transition from subclinical to clinical Cushing’s syndrome. We also test the dynamical function of the pituitary by comparison to alternative designs for the HPA axis, to show that the natural design provides speedup on the scale of weeks to the response to chronic stress.

Hormones are diverse in terms of their biochemistry. The cells that secrete them also differ in many respects, and the size of the secretory glands ranges from a few thousand cells to about 10 billion cells. Despite these differences, we find an approximate proportionality between endocrine gland mass and the total mass of its target tissues. This suggests that across systems, a single endocrine cell serves a fixed number of target cells, about 1,000. One possibility to understand this proportionality is that there is a maximal hormone production rate per cell and that endocrine glands evolved to be just large enough to serve their targets, but not larger. It is interesting for future work to explore these topics.

Each of the five classes of circuits defined here has a distinct regulatory role. Class 1 and 2 circuits can serve as simple input-output devices. Class 3 circuits can lock a metabolite to a tight range around a specific concentration. An example is control of glucose tightly around 5mM.

Class 4 circuits lock the input hormone level and can offer allostatic control of their output metabolite. An example are intestinal hormones secretin and gastrin that control bicarbonate secretion and stomach acid secretion respectively. Finally class 5 circuits are the most complex and can offer compensation for changing physiological parameters, small toxic tumors and other challenges.

The mass of endocrine glands can expand by hypertrophy or hyperplasia. It can also grow in principle by differentiation of progenitor cells. One basic function of such mass growth occurs when high levels of hormones are needed for long times. The circuits of classes 3-5 then sense this need and signal the endocrine cells to grow in total mass. A well known example is endemic goiter in which the thyroid can expand when iodine, essential for thyroid hormone production, is very low(Triggiani *et al*, 2009). The thyroid can grow by a factor of ten or more. The thyroid also grows in pregnancy to meet the needs of the fetus(Gaberšček & Zaletel, 2011). Similarly, the adrenal cortex grows in people under chronic stress or depression(Ulrich-Lai *et al*, 2006; Rubin *et al*, 1995).

Another consequence of the gland mass regulation by these circuits is that they offer a solution to the problem of exponential cell growth. Since cells expand exponentially, their growth and removal rates need to be precisely matched to avoid excess mass growth or shrinkage. The circuits make sure that the gland mass production and removal rates balance precisely when the signal reaches a functional level(Karin *et al*, 2016). Thus, the same circuits solve two problems: expansion of gland mass when more hormone is needed, and organ size control.

One interesting question in class 5 circuits is the purpose of a middle gland-the pituitary in HP axes. Why wouldn’t an alternative design without a middle gland, like class 4 circuits, be chosen by natural selection instead? The findings here propose several answers. First, the middle gland acts as an amplifier. In the HP axes, a small brain region in the hypothalamus secretes a hormone, and the effector gland needs to serve the entire body, and is thus on the order of 10^10^ cells as we found in our mass law. In order to supply such a large effector gland, it is useful to have a middle gland (e.g. 10^7^− 10^8^ for each secretory cell type) so that the hypothalamic region can be small. A tiny amount of hypothalamic hormone thus regulates a sub-pea-sized pituitary, which regulates a large effector gland. In the HP axes, there is about a thousand fold ratio between the top and middle gland, and a similar ratio between the middle and effector gland.

A second functional advantage of the middle gland is speedup of responses. Alternative designs without a middle gland, or with a middle gland that can’t change its mass, have slower response on the scale of weeks. The middle gland in class 5 circuits achieves speedup by causing a mild overshoot or undershoot of several weeks in the hormone dynamics. This overshoot can cause mild dysregulation when entering or exiting prolonged periods of high excitation, as in prolonged stress(Karin *et al*, 2020). Finally, the middle gland can participate in compensation of physiological or pathological perturbations. We demonstrate this by analyzing the effects of hormone-secreting tumors in the HPA axis. A cortisol-secreting tumor in the adrenal can be fully compensated by reduction of the pituitary ACTH-secreting cell mass, provided that the tumor secretion rate is below a threshold value. This avoids clinical consequences of mildly cortisol-secreting tumors, which may account for 15% of incidental tumors (incidentalomas) found in the adrenal(Fassnacht *et al*, 2016). These mildly secreting tumors are thus quite common subclinical events given that incidentalomas are found in about 4% of individuals undergoing high-resolution abdominal imaging(Bovio *et al*, 2006). When the tumor secretion rate crosses a threshold value, however, the effective pituitary mass shrinks to zero, and compensation cannot continue. Thereafter, high cortisol with clinical symptoms can result, a condition known as Cushing’s syndrome.

It would be interesting to compare these results in humans with other organisms. The major hormone regulatory circuits tend to be conserved in mammals and vertebrates, with the main differences being switches between the dominant chemical form of the hormone (e.g. cortisol in humans, corticosterone in mice) and sometimes in its biological functions on target tissues. It is thus plausible that the same five circuit classes will occur across vertebrates.

The question of gland mass ratios in other species also requires further research. Recent advances signal the availability of large scale data in other species in the near future, such as a study on the hormone network of a primate, the mouse lemur(Mouse lemur transcriptomic atlas elucidates primate genes, physiology, disease, and evolution | bioRxiv).

In summary, endocrine systems are diverse in biochemistry, structure, size and physiological function. Despite this diversity, design principles that unite these systems can be found. Gland mass is proportional to the total mass of its target suggesting a fixed capacity of production per endocrine cell. Endocrine regulatory mechanisms fall into 5 classes of recurring circuit motifs with different dynamical functions. These circuits can cause gland masses to grow or shrink to compensate for physiological and pathological changes. It would be interesting to explore whether other design principles can be found to deepen our understanding of systems endocrinology.

## Methods

Analysis of endocrine systems for Figure 1: Of all human hormones(Melmed *et al*, 2019; UpToDate. 2023. Waltham, MA: UpToDate Inc.; [cited 2023 March 21]. Available from: https://www.uptodate.com/home), we chose those with a known cell of origin. We only consider hormones made by dedicated cells whose major function is to produce and secrete the hormone. We do not include hormones made by cells which have a different major function, such as fat or liver cells. These excluded hormones include adiponectin, and leptin made by adipocytes and thrombopoietin, hepcidin, IGF1 made in the liver.

We evaluated the number of secreting cells using a literature search (SI). We evaluated target cells for each hormone using the following criteria: (1) cells that express the receptor for the hormone at high level, based on literature search, (2) cells where the hormone has a documented physiological function, (3) we excluded cases where the receptor is expressed but no physiological function is known. This includes cases where the hormone has only a known pathophysiological effect-for example TSH receptors are found in the eye orbital muscles, but have a known effect only in cases of hyperthyroidism such as Graves disease. We did not include hormones for which we could not find reliable values for the size of the secreting tissue including oxytocin, GHRH, vasopressin, GLP, Ghrelin and motilin. The supplementary information includes a detailed account of the evaluation for each hormone.

Mathematical modeling: We used a minimal model approach inspired by work on insulin that led to the HOMA formulas. We used linear removal terms, and 1/x inhibition terms (which model Michaelis-menten forms 1/(k+x) when x is larger than k). Cell total mass equations used linear dependence on the growth factor and linear dependence on cell mass in the growth term and removal terms linear in the cell mass. We relied on models for the HPA axis(Karin *et al*, 2020) as a basis for our approach.

## Acknowledgements

We thank R. Sender and members of the Alon lab for discussions and comments on the manuscript. Funding was provided by the European Research Council (ERC) under the European Union’s Horizon 2020 research and innovation program (grant agreement No. 856587). D.S.G. was funded as a member of the Zuckerman Postdoctoral Scholars Program. U.A is the incumbent of the Abisch-Frenkel Professional Chair. Yael Korem Kohanim is supported by the JSMF Postdoctoral Fellowship in Understanding Dynamic and Multi-scale Systems (Award #https://doi.org/10.37717/2020-1428).

